# Novel insights into L-serine exporters in *Corynebacterium glutamicum* from gene mining and functional analysis

**DOI:** 10.1101/2022.07.13.499997

**Authors:** Yujie Gao, Xiaomei Zhang, Guoqiang Xu, Xiaojuan Zhang, Hui Li, Jinsong Shi, Zhenghong Xu

## Abstract

Amino acid exporters play an important role in regulating amino acid production by *Corynebacterium glutamicum*, and over 90% of amino acid export is attributed to exporters in this species. ThrE was reported to be an l-serine exporter, and SerE was identified as an l-serine exporter in our previous study. However, when both ThrE and SerE were deleted, the l-serine titer was decreased by 60%, suggesting other l-serine exporters may exist. In the present study, NCgl0254 and NCgl0255 were identified as novel l-serine exporters through comparative transcriptomics and gene functional analyses. The contributions of the four exporters (ThrE, SerE, NCgl0254 and NCgl0255) in l-serine export were studied by gene deletion, gene overexpression, amino acid export assay and real-time quantitative PCR (RT-qPCR). The results showed that SerE is the major l-serine exporter in *C. glutamicum*. Fermentation and amino acid export assays of SSAAI, SSAAI-*serE*-*thrE*-*ncgl0254*-*ncgl0255* and SSAAIΔ*serE*Δ*thrE*Δ*ncgl0254*Δ*ncgl0255* indicated that the four l-serine exporters undertake most of the l-serine export, and their overexpression enhanced export of l-serine in SSAAI. When one l-serine exporter was deleted, the transcription level of the other three exporters was upregulated. However, the decrease in l-serine titer caused by deletion of one exporter was not fully compensated by upregulation of the other three exporters at the transcription level, indicating that l-serine production by *C. glutamicum* may be determined by cooperative efficiency of all four l-serine exporters, with each being interdependent.

**IMPORTANCE:** This work identified the novel l-serine exporters NCgl0254 and NCgl0255, and revealed their roles in l-serine export alongside the l-serine exporters. All four l-serine exporters are interdependent and undertake most of the l-serine export, but SerE is the major l-serine exporter. The findings expand our knowledge of amino acid exporters in *C. glutamicum,* and the approach can be employed for exploring of bacterial exporters of unknown function.

## INTRODUCTION

l-serine is a biochemical building block with potential as a neuroprotective agent and a drug for the treatment of nervous system diseases (1, 2). *Corynebacterium glutamicum* is widely used for producing l-serine by fermentation due to its high safety and clear genetic background (3, 4). Metabolic engineering of *C. glutamicum* for l-serine production has focused on synthesis and degradation pathways, when l-serine pathway genes were overexpressed, a feedback-resistant variant was obtained, degradation of l-serine to pyruvate was avoided by deletion of *sdaA*, and downregulation of *glyA* further enhanced l-serine production (4–8). However, little attention has been paid to l-serine export. Amino acid exporters play an important role in regulating amino acid production by *C. glutamicum*, and over 90% of amino acid export is attributed to exporters in this species (9, 10).

In the past few years, only seven amino acid exporters have been identified in *C. glutamicum*, involved in the export of 13 amino acids (Fig. 1) (11–20). Among the seven exporters, BrnFE participates in the export of l-methionine, l-isoleucine, l-valine, and l-leucine (11, 12). CgmA participates in the export of l-arginine (13). LysE participates in the export of l-lysine, l-arginine, l-citrulline, and l-ornithine (13, 14). MscCG participates in the export of l-glutamic and l-aspartic (15, 16). MscCG2 participates in the export of l-glutamic (17). ThrE was first identified as an l-threonine exporter in *C. glutamicum*. Simic et al. isolated l-threonine resistance mutants, and identified the l-threonine exporter ThrE by analysing phenotypic and genotypic changes between mutant and wild-type strains. Since the chemical structure of l-serine is similar to that of l-threonine, ThrE can export l-serine (18, 21). Recent studies by arrayed CRISPRi screening showed that ThrE is also an exporter of l-proline (20). In our previous studies, we found that overexpressing or deleting *thrE* in SSAAI (with an l-serine titer of 26 g/L) had no significant effect on l-serine production. Thus, other l-serine exporters were suspected, and the novel l-serine and l-threonine exporter SerE was identified in *C. glutamicum* by homologous sequence alignment with EamA in *Escherichia coli* (19, 22). However, 40% of l-serine was exported out of the cell when both *thrE* and *serE* were deleted. This indicates that other exporters are involved in l-serine export in *C. glutamicum*. Thus, it is important to further explore amino acid exporters, clarify their relationships, and thereby improve amino acid export efficiency.

**FIG 1.**
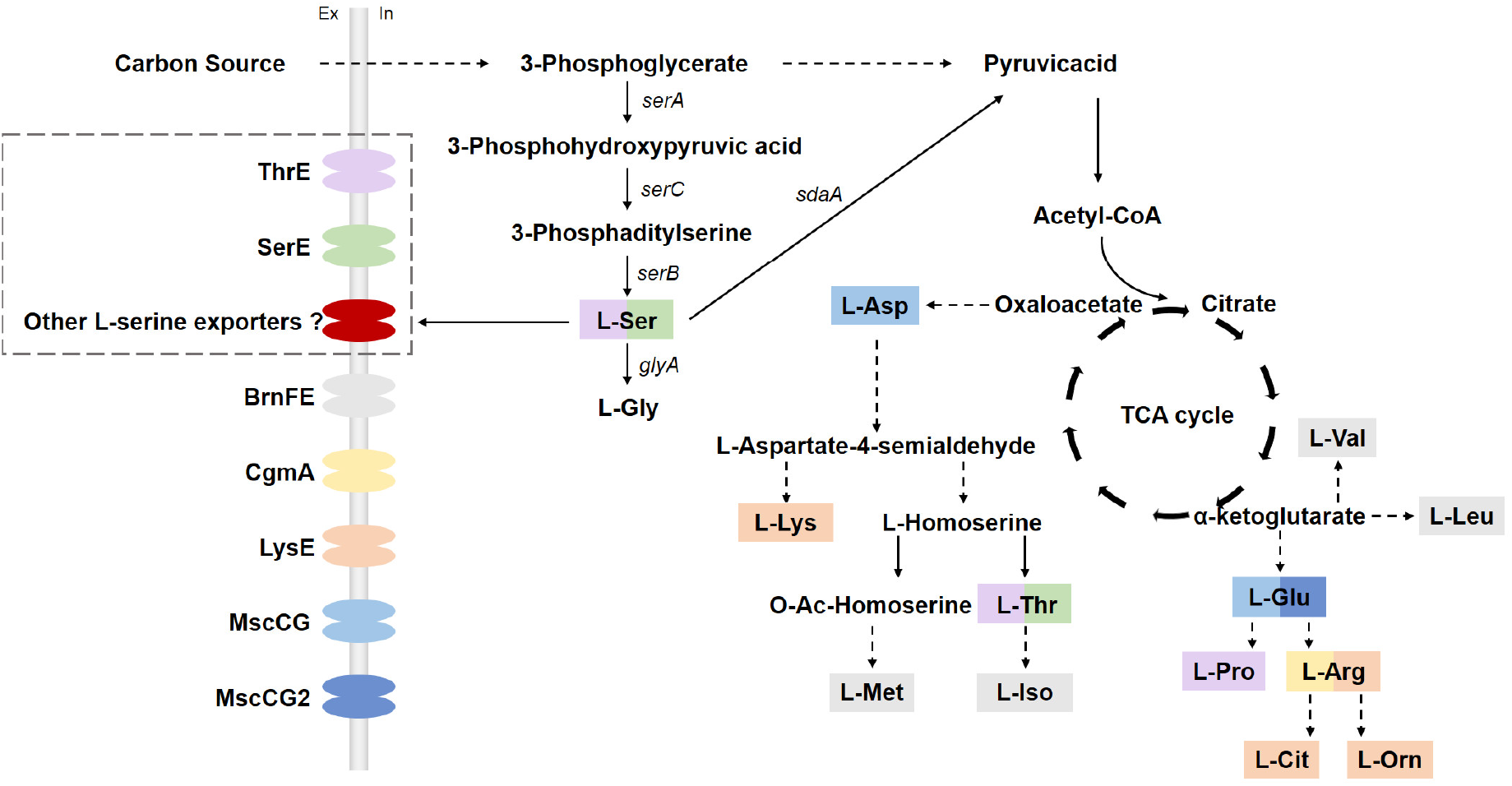
Schematic representation of amino acid exporters in *C. glutamicum*. ThrE (purple) exports l-threonine, l-serine and l-proline. SerE (green) export l-threonine and l-serine. BrnFE (grey) exports l-valine, l-leucine, l-isoleucine and l-methionine. CgmA (yellow) exports l-arginine. LysE (orange) exports l-lysine, l-arginine, l-citrulline and l-ornithine. MscCG (blue) exports l-glutamate and l-aspartate. MscCG2 (dark blue) exports l-glutamate.

In the present study, l-serine exporters were identified via comparative transcriptomics and differential gene functional analyses of four strains with different l-serine titers (SYPS-062, 6.6 g/L; SYPS-062-33a, 13.2 g/L; SSAAI, 26 g/L; A36, 31 g/L) (19, 23–26). Next, the contribution of each exporter to l-serine export was studied by gene deletion, gene overexpression and amino acid export assay. Finally, the relationships of l-serine exporters SerE, ThrE, and the novel exporters identified in this study were investigated by RT-qPCR. This study explored novel amino acid exporters and clarified their relationships, and the findings provide novel insight into efficient amino acid export.

## RESULTS

### Mining genes related to l-serine export via comparative transcriptomics

In general, when a strain produces a high titer of amino acids, the expression level of the corresponding exporter is usually high (27). To mine novel l-serine exporters, we comparatively analysed transcripts extracted from four strains with different l-serine titer (SYPS-062, SYPS-062-33a, SSAAI and A36) at the logarithmic phase of fermentation (60 h). The genes showed more than a four-fold greater expression level in high-yielding strains, and they were annotated as encoding membrane proteins or hypothetical proteins by comparing results of gene expression level (transcripts per million), Gene Ontology (GO), and Kyoto Encyclopedia of Genes and Genomes (KEGG) analyses, with log_2_ Fold Change >2 and *q* <0.05 as criteria for pairwise comparisons. The pairwise comparison groups included SYPS-062-33a vs. SYPS-062 (VS1), SSAAI vs. SYPS-062 (VS2), SSAAI vs. SYPS-062-33a (VS3), A36 vs. SYPS-062 (VS4), A36 vs. SYPS-062-33a (VS5) and A36 vs. SSAAI (VS6), and the results are shown in Fig. 2A.

**FIG 2.**
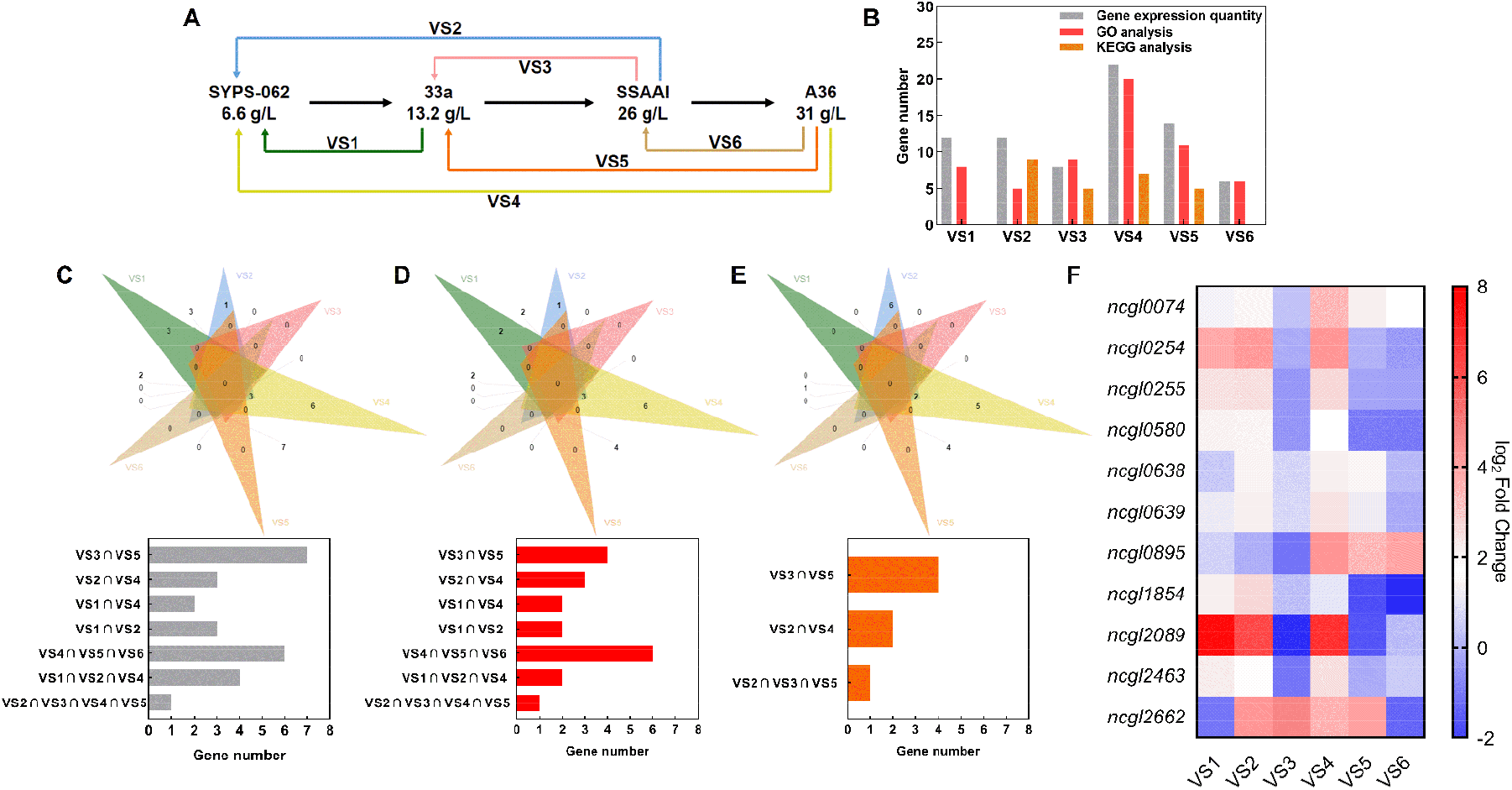
Transcriptome analysis of four strains (SYPS-062, SYPS-062-33a, SSAAI and A36) with different l-serine titer. (A) Pairwise comparison groups for the four strains. VS1, SYPS-062-33a vs. SYPS-062; VS2, SSAAI vs. SYPS-062; VS3, SSAAI vs. SYPS-062-33a; VS4, A36 vs. SYPS-062; VS5, A36 vs. SYPS-062-33a; VS6, A36 vs. SSAAI. (B) Number of genes in pairwise comparison groups for gene expression level, GO analysis and KEGG analyses. (C) Venn diagram of differential genes from pairwise comparison of gene expression. (D) Venn diagram of differential genes from GO analysis. (E) Venn diagram of differential genes from KEGG analysis. (F) Transcriptional changes for genes encoding potential l-serine exporters in pairwise comparison groups.

Pairwise comparison of gene expression levels, GO terms and KEGG pathways for the four strains are shown in Fig. 2B. Venn diagrams were used to analyse upregulated genes among the pairwise comparison groups. The number of genes upregulated in more than two groups for gene expression level, GO term and KEGG pathway results was 26, 20 and 7, respectively (Fig. 2C−E). By combining the three Venn analyses, removing repeated genes, and searching functions in the NCBI database, 11 genes were identified, of which *ncgl0255* and *ncgl0254* were annotated as branched-chain amino acid transporters, *ncgl2089*, *ncgl2662* and *ncgl1854* as hypothetical proteins with unknown functions, *ncgl2463* as related to amino acid transport, and *ncgl0639*, *ncgl0074*, *ncgl0895*, *ncgl0638* and *ncgl0580* as transporters. The fold changes in expression for the 11 genes are shown in Fig. 2F. The *ncgl0074*, *ncgl0580*, *ncgl0638*, *ncgl0639*, *ncgl1854* and *ncgl2463* genes were upregulated in two pairwise comparison groups, *ncgl0254*, *ncgl0255*, *ncgl0895* and *ncgl2089* were upregulated in three pairwise comparison groups, and *ncgl2662* was upregulated in four pairwise comparison groups. These results indicate that the 11 genes might encode potential l-serine exporters, hence their functions were further investigated.

### Verification of the functions of potential l-serine exporters

In order to verify the functions of the 11 genes, two rounds of verification were conducted. Firstly, since deletion of l-serine exporters can significantly decrease the l-serine titer, the 11 genes were separately deleted in strain SSAAI, and the resulting recombinant strains were obtained and tested in fermentation experiments (Table 1). The fermentation results showed that the l-serine titer of SSAAIΔ*ncgl0638*, SSAAIΔ*ncgl0639*, SSAAIΔ*ncgl0254*, SSAAIΔ*ncgl0255* and SSAAIΔ*ncgl0580* were decreased by 10%, 11%, 17.2%, 18.6% and 56.5%, respectively, compared with the parent strain SSAAI, while the l-serine titers of other strains were similar to that of SSAAI. NCgl0580 was identified as l-serine exporter SerE in our previous study (19). After deleting *ncgl0638* and *ncgl0254* in SSAAI, the growth (OD_562_) of strains showed a similar trend to that of SSAAI. Also, the growth (OD_562_) of SSAAI strains lacking *ncgl0639* or *ncgl0255* displayed a similar trend to strain SSAAI before 84 h, but after 96 h the growth of these deletion strains was faster than that of the parent strain (Fig. 3A). As shown in Fig. 3B, deleting each of the four genes (*ncgl0638*, *ncgl0639*, *ncgl0254* and *ncgl0255*) individually had an impact on the accumulation of l-serine at the early stages of growth. At 12 h and 24 h of fermentation, the amount of l-serine in all deletion strains was significantly lower than that in strain SSAAI, and eventually led to a decrease of more than 10% in l-serine titer at 120 h. These results suggest that the four proteins encoded by *ncgl0638*, *ncgl0639*, *ncgl0254* and *ncgl0255* might be l-serine exporters.

**FIG 3.**
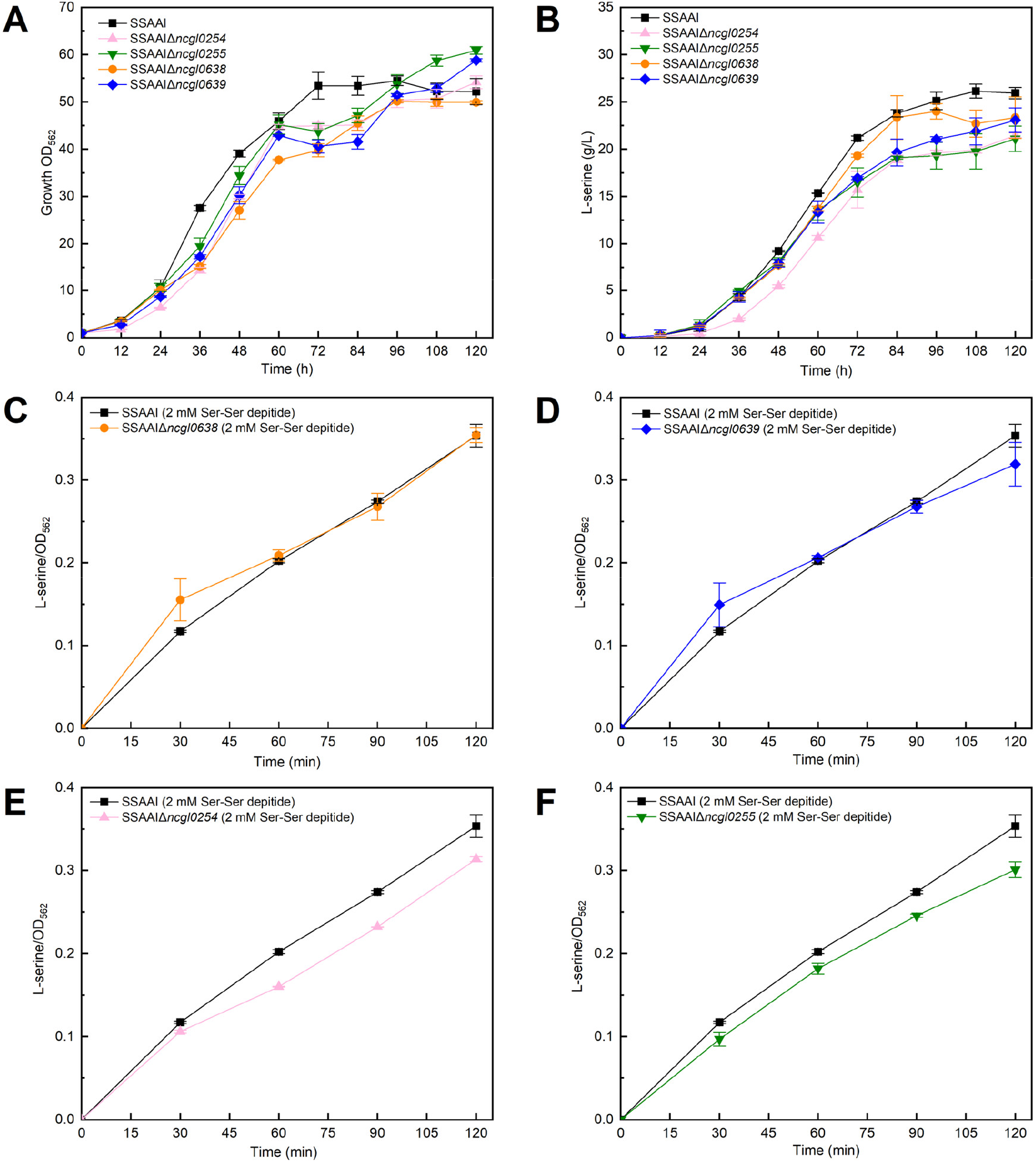
Verification of l-serine exporter functions. (A) Cell growth (OD_562_) of SSAAIΔ*ncgl0254*, SSAAIΔ*ncgl0255*, SSAAIΔ*ncgl0638* and SSAAIΔ*ncgl0639*. (B) l-serine titer of SSAAIΔ*ncgl0254*, SSAAIΔ*ncgl0255*, SSAAIΔ*ncgl0638* and SSAAIΔ*ncgl0639*. (C−F) Amino acid export assays for SSAAI, SSAAIΔ*ncgl0254*, SSAAIΔ*ncgl0255*, SSAAIΔ*ncgl0638* and SSAAIΔ*ncgl0639*. Three parallel experiments were performed. Error bars indicate standard deviations of results from three parallel experiments.

**TABLE 1.**
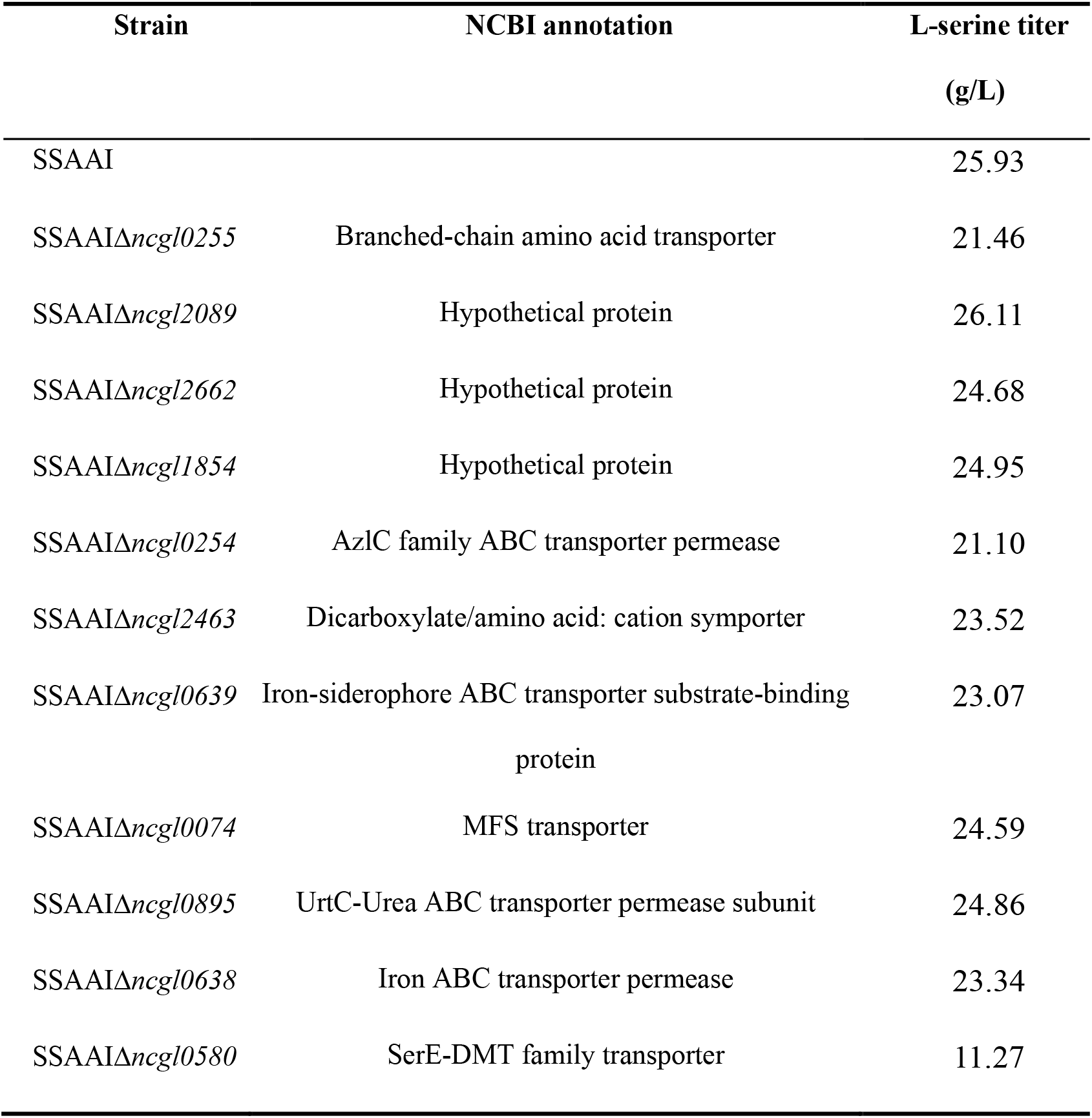
NCBI annotation of genes and l-serine titers of deletion strains

Next, to further explore the functions of NCgl0638, NCgl0639, NCgl0254 and NCgl0255, classical amino acid dipeptide export assays were conducted, which have been widely employed to verify amino acid exporters in *C. glutamicum* (18). Upon addition of 2 mM Ser-Ser peptide, the l-serine yield to biomass ratio of SSAAIΔ*ncgl0638* and SSAAIΔ*ncgl0639* was no different to that of SSAAI (Fig. 3C and Fig. 3D). These results suggest that NCgl0638 and NCgl0639 are unrelated to l-serine export. As shown in Fig. 3E, upon addition of 2 mM Ser-Ser peptide, the l-serine yield to biomass ratio of SSAAIΔ*ncgl0254* throughout the entire export process was lower than that of SSAAI, and decreased by 11.3% compared with SSAAI at 120 min. These results confirmed that NCgl0254 is a novel l-serine exporter. Upon addition of 2 mM Ser-Ser peptide, the l-serine yield to biomass ratio of SSAAIΔ*ncgl0255* was lower than that of SSAAI, and decreased by 14.9% compared with SSAAI at 120 min (Fig. 3F). These results indicate that NCgl0255 is another novel l-serine exporter.

### Contributions of the four exporters to l-serine export

SerE and ThrE have been reported as l-serine exporters, while NCgl0254 and NCgl0255 were confirmed to be novel l-serine exporters in the above experiments. However, the contributions of the four exporters (ThrE, SerE, NCgl0254 and NCgl0255) to l-serine export in *C. glutamicum* remained unclear. Hence, we first examined the effect of deleting each gene (*thrE*, *serE*, *ncgl0254* and *ncgl0255*) on l-serine production by *C. glutamicum*. As shown in Fig. 4A, compared with SSAAI (S1), the l-serine titer of SSAAIΔ*serE* (S2), SSAAIΔ*thrE* (S3), SSAAIΔ*ncgl0254* (S4) and SSAAIΔ*ncgl0255* (S5) was decreased by 56.5%, 3.3%, 18.6% and 17.2%, respectively. There was no significant change in growth (OD_562_) for any of the single deletion strains compared with SSAAI at 120 h. However, a significant decrease in l-serine accumulation was observed for SSAAIΔ*serE* (S2), while only a slight decrease was observed for SSAAIΔ*thrE* (S3). Moreover, the l-serine yield to biomass (Y_p/x_) of SSAAIΔ*serE* (S2) was decreased by 58.4% compared with SSAAI (S1). These results indicate that SerE might be the major l-serine exporter in *C. glutamicum*.

**FIG 4.**
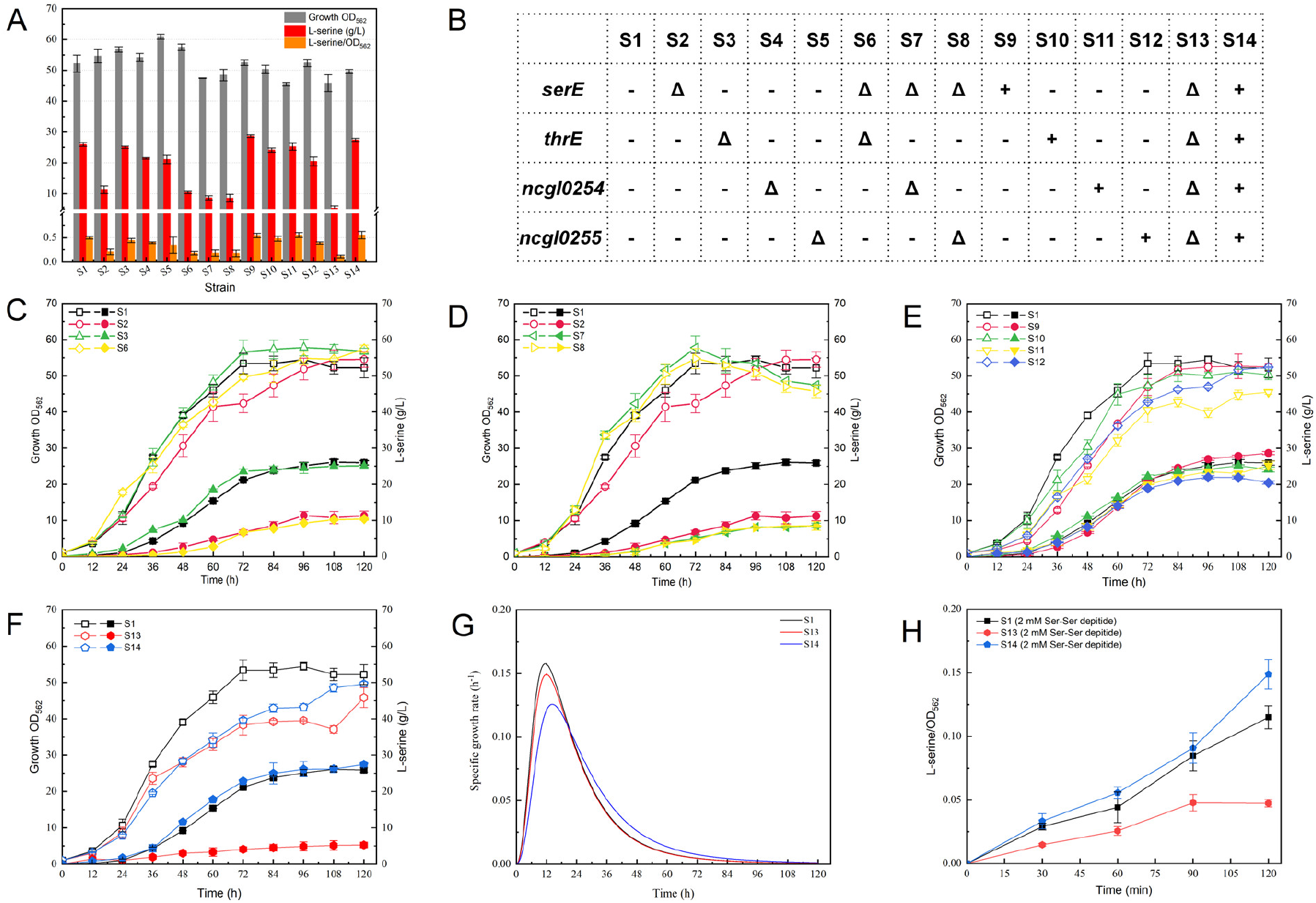
Effects of *serE*, *thrE*, *ncgl0254* and *ncgl0255* deletion on SSAAI. (A) Cell growth (OD_562_) (grey), l-serine titer (red) and Y_P/X_ (orange) of SSAAI, single deletion, and multiple deletion strains. S1, SSAAI; S2, SSAAIΔ*serE*; S3, SSAAIΔ*thrE*; S4, SSAAIΔ*ncgl0254*; S5, SSAAIΔ*ncgl0255*; S6, SSAAIΔ*serE*Δ*thrE*; S7, SSAAIΔ*serE*Δ*ncgl0254*; S8, SSAAIΔ*serE*Δ*ncgl0255*; S9, SSAAI-*serE*; S10, SSAAI-*thrE;* S11, SSAAI-*ncgl0254;* S12, SSAAI-*ncgl0255;* S13, SSAAIΔ*serE*Δ*thrE*Δ*ncgl0254*Δ*ncgl0255;* S14, SSAAI-*serE*-*thrE*-*ncgl0254*-*ncgl0255*. (B) Gene editing details of strains: Δ represents gene deletion; + represents gene overexpression;-represents no editing of the gene. (C) Effects of *serE* and *thrE* deletion on SSAAI. The hollow pattern represents cell growth (OD_562_) and the solid pattern represents l-serine titer. (D) Effects of *serE*, *ncgl0254* and *ncgl0255* deletion on SSAAI. The hollow pattern represents cell growth (OD_562_) and the solid pattern represents l-serine titer. (E) Effects of *serE*, ThrE, *ncgl0254* and *ncgl0255* overexpression on SSAAI. The hollow pattern represents cell growth (OD_562_) and the solid pattern represents l-serine titer. (F) Effects of combined *serE*, *ncgl0254* and *ncgl0255* deletion or overexpression on SSAAI. The hollow pattern represents cell growth (OD_562_) and the solid pattern represents l-serine titer. (G) The growth rates of SSAAIΔ*serE*Δ*thrE*Δ*ncgl0254*Δ*ncgl0255* and SSAAI-*serE*-*thrE*-*ncgl0254*-*ncgl0255*. (H) Amino acid export assays for SSAAI, SSAAIΔ*serE*Δ*thrE*Δ*ncgl0254*Δ*ncgl0255* and SSAAI-*serE*-*thrE*-*ncgl0254*-*ncgl0255*. Three parallel experiments were performed. Error bars indicate standard deviations of results from three parallel experiments.

To further explore the contributions to l-serine production of the four l-serine exporters, *thrE*, *ncgl0254* and *ncgl0255* were each separately deleted alongside *serE*, resulting in strains SSAAIΔ*serE*Δ*thrE* (S6), SSAAIΔ*serE*Δ*ncgl0254* (S7) and SSAAIΔ*serE*Δ*ncgl0255* (S8). As shown in Fig. 4C, the l-serine titer of SSAAIΔ*serE*Δ*thrE* (S6) was decreased by 60% compared with SSAAI (S1), and decreased by 8% compared with SSAAIΔ*serE* (S2). Obviously, a major proportion of the decrease in the l-serine titer in SSAAIΔ*serE*Δ*thrE* (S6) was attributed to the deletion of *serE*. However, when *thrE* was deleted together with *serE*, the strain could still export l-serine (Fig. 4C), and a slight decrease in cell growth was observed compared with SSAAIΔ*serE* (S2). As shown in Fig. 4D, the l-serine titer of SSAAIΔ*serE*Δ*ncgl0254* (S7) and SSAAIΔ*serE*Δ*ncgl0255* (S8) was decreased by 66.7% and 66.1%, respectively, compared with SSAAI (S1), and decreased by 23.4% and 24.4% compared with SSAAIΔ*serE* (S2). As shown in Fig. 4C, compared with other combinations of deletion strains, when *ncgl0254* or *ncgl0255* were deleted with *serE*, the resulting strains exhibited higher growth rates than SSAAI (S1) and SSAAIΔ*serE* (S2) before 72 h, but the cell growth of the deletion strains was decreased significantly after 72 h, and decreased by 13% and 12% at 120 h, compared with SSAAIΔ*serE* (S2). From the fermentation results for single and multiple gene deletion strains, deletion of *serE* appears to be the main reason for the diminished l-serine titer. Thus, SerE is the major l-serine exporter, followed by NCgl0254 and NCgl0255, with ThrE the least important.

Next, *serE*, *thrE*, *ncgl0254* and *ncgl0255* were overexpressed in SSAAI to obtain strains SSAAI-*serE* (S9), SSAAI-*thrE* (S10), SSAAI-*ncgl0254* (S11) and SSAAI-*ncgl0255* (S12), respectively. As shown in Fig. 4E, a decrease in the growth (OD_562_) of the overexpression strains was observed before 108 h of fermentation compared with SSAAI (S1), especially SSAAI*-ncgl0254* (S11) with a decrease of >12.8% compared with SSAAI (S1). As shown in Fig. 4E, the l-serine titer of SSAAI-*serE* (S9) was 10.6% higher than that of SSAAI (S1). However, the l-serine titers of SSAAI-*thrE* (S10) and SSAAI-*ncgl0254* (S11) were similar to that of SSAAI (S1). As shown in Fig. 4A, the Y_p/x_ of SSAAI-*ncgl0254* (S11) was 11.7% higher than that of SSAAI (S1), and the Y_p/x_ of SSAAI-*serE* (S9) was 9.9% higher than that of SSAAI (S1). The results indicate that overexpression of l-serine exporters contributed to l-serine export.

We then engineered SSAAIΔ*serE*Δ*thrE*Δ*ncgl0254*Δ*ncgl0255* (S13) and SSAAI-*serE*-*thrE*-*ncgl0254*-*ncgl0255* (S14), overexpressing or deleting the four l-serine exporters, to further explore their contributions. As shown in Fig. 4A and 4F, the l-serine titer of SSAAIΔ*serE*Δ*thrE*Δ*ncgl0254*Δ*ncgl0255* (S13) was decreased by 79.7% and Y_p/x_ was decreased by 76.9% compared with SSAAI (S1). Since more than 90% of amino acid secretion in *C. glutamicum* is undertaken by exporters, the four exporters re clearly responsible for most of the l-serine export in this species (9, 10). As shown in Fig. 4A and 4F, the l-serine titer of SSAAI-*serE*-*thrE*-*ncgl0254*-*ncgl0255* (S14) was similar to that of SSAAI (S1), but Y_p/x_ was 11.3% higher than for SSAAI (S1). This indicates that overexpression of exporters is beneficial for l-serine efflux, but to improve the l-serine titer needs the relationship between growth and l-serine accumulation must be balanced. As shown in Fig. 4F, the growth (OD_562_) of SSAAIΔ*serE*Δ*thrE*Δ*ncgl0254*Δ*ncgl0255* (S13) and SSAAI-*serE*-*thrE*-*ncgl0254*-*ncgl0255* (S14) was 11.3% and 7.7% lower than for SSAAI at 120 h. As shown in Fig. 4G, the growth rates of SSAAIΔ*serE*Δ*thrE*Δ*ncgl0254*Δ*ncgl0255* (S13) and SSAAI-*serE*-*thrE*-*ncgl0254*-*ncgl0255* (S14) were lower than for SSAAI (S1), especially SSAAI-*serE*-*thrE*-*ncgl0254*-*ncgl0255* (S14). This might be due to inhibition of cell growth resulting excessive l-serine efflux or intracellular accumulation. To further explore the contributions of overexpression or deletion of the four l-serine exporter-encoding genes in SSAAI to l-serine export, amino acid export assays were conducted. As shown in in Fig. 4H, the l-serine yield to biomass ratio of SSAAIΔ*serE*Δ*thrE*Δ*ncgl0254*Δ*ncgl0255* (S13) was decreased by 58.8% compared with SSAAI (S1), implying that the four l-serine exporters undertake most of the l-serine export. The l-serine yield to biomass ratio of SSAAI-*serE*-*thrE*-*ncgl0254*-*ncgl0255* (S14) was increased by 29.4% compared with SSAAI, which suggests that overexpression of the four exporters enhanced the l-serine export ability of SSAAI.

### Interactions of the four l-serine exporters during the fermentation process

SerE was identified as the major l-serine exporter, and the contributions of the four l-serine exporters were determined in the above studies. However, the relationships between the four l-serine exporters remained unknown. Expression levels of *serE*, *thrE*, *ncgl0254* and *ncgl0255* in four different phenotypic strains (SYPS-062, STPS-062-33a, SSAAI and A36) at 60 h were analysed by transcriptome sequencing. First, expression levels of the exporters in the four different strains were compared, and the results are shown in Fig. 5A. Compared with SYPS-062 (l-serine titer 6.6 g/L), expression levels of *serE*, *thrE*, *ncgl0254* and *ncgl0255* in SYPS-062-33a (l-serine titer 13.2 g/L) were increased by 3.69, 1.29, 5.6 and 14.5 times, respectively. Compared with SYPS-062-33a, the expression levels of *ncgl0254* and *ncgl0255* in SSAAI (l-serine titer 26 g/L) were increased by 1.13 and 1.27 times, respectively, whereas there was no significant change in *serE* and *thrE*. Compared with SSAAI, the expression levels of *ncgl0254* and *ncgl0255* in A36 (l-serine titer 31 g/L) were increased by 1.13 and 1.07 times, respectively. Interestingly, there was no obvious change in the expression levels of *serE* or *thrE*. This indicates that an increase in the expression of exporters is very important for enhancing l-serine production. As shown in Fig. 5A, compared with the major l-serine exporter SerE, the expression levels of the other three exporters in the four different strains were relatively lower. Expression levels of *thrE*, *ncgl0254* and *ncgl0255* were only 6.8%, 2.4% and 2.3% those of *serE* in SYPS-062 (l-serine titer 6.6 g/L), respectively. Expression levels of *thrE*, *ncgl0254* and *ncgl0255* were only 2.4%, 9.7% and 3.4% those of *serE* in strain SYPS-062-33a (l-serine titer 13.2 g/L), respectively. Expression levels of *thrE*, *ncgl0254* and *ncgl0255* were only 2.3%, 12.1% and 3.5% those of *serE* in strain SSAAI (l-serine titer 26 g/L), respectively. Expression levels of *thrE*, *ncgl0254* and *ncgl0255* were only 1.7%, 16.6% and 5.1% those of *serE* in strain A36 (l-serine titer 31 g/L), respectively. Expression levels of *serE* were consistently the highest among the four l-serine exporters, and expression levels of the other three l-serine exporters were relatively lower. These results further indicate that SerE is the major l-serine exporter in *C. glutamicum*, explaining why deletion of *serE* had a significant effect on l-serine production. Combining the two analyses, we found that the expression levels of all l-serine exporters were higher with increasing l-serine titer, and the expression level of *serE* was consistently the highest. These results indicate that all four l-serine exporters contribute to determining the l-serine titer, hence it is useful to investigate the relationships among the four l-serine exporters.

**FIG 5.**
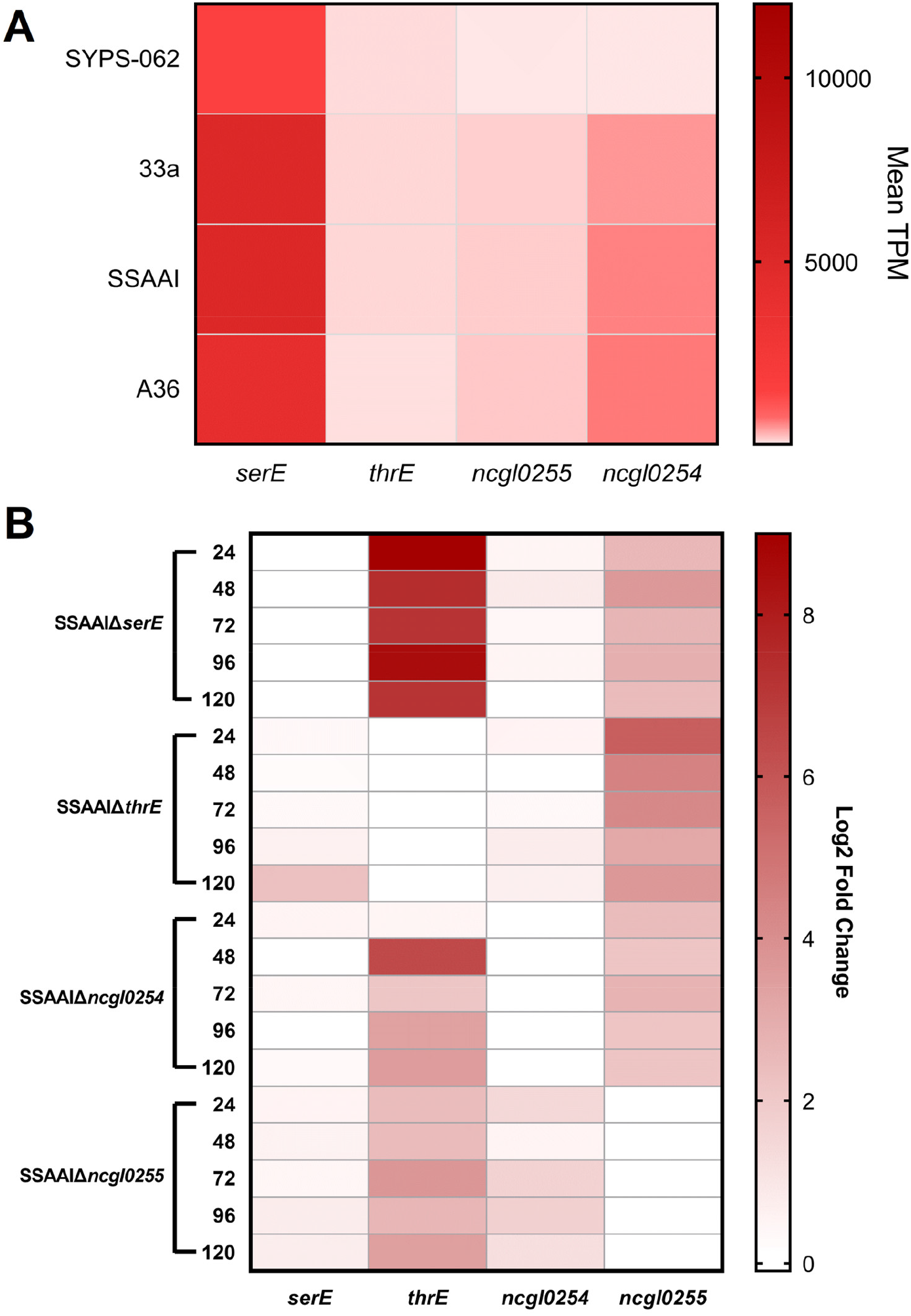
Expression and transcription levels of the four exporters. (A) Expression levels of the four exporters in four strains with different phenotypes. (B) Transcription level changes of the four exporters in single gene deletion strains. Three parallel experiments were performed. Error bars indicate standard deviations of results from three parallel experiments.

To further explore the interactions of the four l-serine exporters, we analysed changes in the transcription levels of the other three exporters when one was deleted. To this end, changes in the transcription levels of the four genes (*serE*, *thrE*, *ncgl0254* and *ncgl0255*) in SSAAIΔ*serE*, SSAAIΔ*thrE*, SSAAIΔ*ncgl0254* and SSAAIΔ*ncgl0255* were measured by RT-qPCR. Transcription level changes (Log_2_ Fold Change) of the four single exporter deletion strains at five time points (24 h, 48 h, 72 h, 96 h and 120 h) were compared with those of SSAAI. As shown in Fig. 5B, when *serE* was deleted, the transcription levels of *thrE* and *ncgl0255* were increased by 143.4 and 5.5 times, respectively. When *thrE* was deleted, the transcription levels of *serE*, *ncgl0254* and *ncgl0255* were increased by 8.7, 1.6 and 12.6 times, respectively. When *ncgl0254* was deleted, the transcription levels of s*erE*, *thrE* and *ncgl0255* were increased by 1.2, 11.7 and 4.4 times, respectively. When *ncgl0255* was deleted, the transcription levels of *serE*, *thrE* and *ncgl0254* were increased by 1.7, 11 and 2.4 times, respectively. These results showed that when one of the l-serine exporters (the target l-serine exporter) was deleted, the transcription levels of the other three exporters were increased to compensate and prevent a dramatic decline in l-serine titer. However, the l-serine titer following fermentation of SSAAIΔ*serE*, SSAAIΔ*thrE*, SSAAIΔ*ncgl0254* and SSAAIΔ*ncgl0255* showed that upregulation of other l-serine exporters could not completely compensate for the loss of the main (target) l-serine exporter.

### Conclusion

In this study, two novel l-serine exporters, NCgl0254 and NCgl0255, were identified and characterised through comparative transcriptomics, gene deletion, and amino acid dipeptide assay analyses. SerE was found to be the major l-serine exporter based on gene deletion and RT-qPCR. Overexpression of the four l-serine exporters (SerE, ThrE, NCgl0254 and NCgl0255) enhanced the l-serine export ability of SSAAI. The four l-serine exporters were confirmed to undertake most of the l-serine export based on deleting each of the encoding genes. l-serine production by *C. glutamicum* is dependent on cooperative efficiency among four interdependent l-serine exporters.

## DISCUSSION

The amino acid metabolism and export ability of *C. glutamicum* are the basis for its high production characteristics (4). Identification and modification of amino acid exporters have attracted significant research interest in recent years (9, 28, 29). However, to date, only seven amino acid exporters (BrnFE, LysE, CgmA, MscCG, MscCG2, ThrE and SerE) have been identified in *C. glutamicum*. Identification and characterisation of amino acid exporters are important not only for understanding the mechanism of hyperproduction of amino acids, but also for engineering the producers.

Mining of amino acid exporters is important for metabolic engineering of amino acid production. The l-threonine exporter ThrE was identified through gene sequence alignment, and since the structures of l-serine and l-threonine are similar, it was unsurprising that ThrE also possessed the ability to export l-serine, even though at a lower efflux rate (18). In our previous study, when *thrE* was deleted or overexpressed, there was no significant change in the l-serine titer, indicating that other l-serine exporters may exist. Three proteins potentially related to l-serine export were selected by homologous alignment of l-serine exporter EamA in *E. coli*, and SerE was proven to be an l-serine exporter in *C. glutamicum* by gene deletion and amino acid export assay (19). When *serE* and *thrE* were deleted in SSAAI, 40% of l-serine was still exported, which also implies other l-serine exporters in *C. glutamicum*.

In the present study, NCgl0254 and NCgl0255 were identified as candidate l-serine exporters by comparison of transcriptomics sequencing, and verified by gene deletion, strain fermentation, and amino acid export assay. NCgl0254 and NCgl0255 have been named as BrnFE and verified as branched-chain amino acids and l-methionine exporters (11, 12). BrnFE was found to export l-isoleucine, l-leucine and l-valine by transposon mutagenesis and gene deletion (11). BrnFE export of l-methionine was assessed by DNA microarray and gene deletion, and the transport rates for l-isoleucine, l-leucine and l-methionine by BrnFE were similar, but the transport rate of l-valine by BrnFE was lower (12). Recent research showed that BrnFE may also possess the ability to export l-homoserine (30). However, whether BrnFE can export l-serine remains unknown. Since the chemical structure of l-serine is similar to that of l-threonine, we explored whether BrnFE could export l-threonine. Amino acid export assays with dipeptides (2 mM Thr-Thr) were performed for strains SSAAI, SSAAIΔ*ncgl0254* and SSAAIΔ*ncgl0255*. The results revealed that the titer of l-threonine did not differ significantly among strains, indicating that NCgl0254 and NCgl0255 might not export l-threonine (data not shown).

Since amino acid exporters are membrane proteins, it is difficult to analyse the crystal structure (31). Currently, there are no structures published for amino acid exporters in *C. glutamicum*, and only a few *E. coli* amino acid exporter structures have been reported (32, 33). Due to the difficulties of structural analysis, understanding the relationships between amino acid exporters at the transcriptional level can provide knowledge that may be pertinent to the mechanism of amino acid export. In *C. glutamicum*, l-glutamic is exported by MscCG and MscCG2, and l-arginine is exported by LysE and CgmA (13, 34). Although one amino acid with different exporters has been reported, the contributions and interactions among exporters have not been studied. In the present study, the contributions of the four l-serine exporters (SerE, ThrE, NCgl0254 and NCgl0255) to l-serine export were studied through combinatorial gene deletion and gene overexpression. Single deletion and double deletion (*thrE*, *ncgl0254* or *ncgl0255* with *serE*) strains were constructed. As shown in Fig. 4A, the growth (OD_562_) of the single deletion strains at lag phase was higher than SSAAI, and the growth (OD_562_) of single deletion strains at the stationary phase was lower than SSAAI. This may be because deletion of l-serine exporters could still meet l-serine demand at lag phase, but accumulation of l-serine in cells is toxic. When *serE* was deleted, the l-serine titer of SSAAIΔ*serE* decreased by 56.5%, and the fermentation of double deletion strains showed that deletion of *serE* was primarily responsible for the decline in l-serine titer. The contributions of l-serine exporters to l-serine export in *C. glutamicum* were found to be ordered SerE > NCgl0254 > NCgl0255 > ThrE. When the four exporters were separately overexpressed, only the l-serine titer of SSAAI-*serE* was increased (by 10.6%), while the l-serine titers of other overexpression strains were similar to that of SSAAI. When the four exporters were overexpressed together, the l-serine titer was increased by 5.8% compared with SSAAI, and growth (OD_562_) was affected. The Y_P/X_ of SSAAI-*serE*-*thrE*-*ncgl0254*-*ncgl0255* was increased by 11.3% compared with SSAAI. The l-serine titer of SSAAI-*serE*-*thrE*-*ncgl0254*-*ncgl0255* was increased by 29.4% compared with SSAAI based on amino acid export assays. Fermentation and amino acid export assay results indicate that overexpression of the four exporters could enhance the l-serine efflux. This also confirmed that overexpression of exporters could increase amino acid production in *C. glutamicum* (13, 20, 35). However, in order to further improve the l-serine titer, cell growth and l-serine accumulation must be balanced, and directed evolution and rational design of amino acid exporters could be employed to enhance l-serine efflux (36). When the four l-serine exporters were deleted together, the l-serine titer of SSAAI-*serE*Δ*thrE*Δ*ncgl0254*Δ*ncgl0255* was decreased by 79.7% compared with SSAAI, which indicates that the four l-serine exporters undertake most of the l-serine export, but there remained other ways to export l-serine, such as active efflux (9, 10).

Examining transcription levels of key genes is often used to analyse the relationships among proteins (37, 38). Transcriptomics sequencing analysis of the four strains with different l-serine titers was subsequently performed, and the results further indicated that SerE is the major l-serine exporter in *C. glutamicum*. Transcription levels of *serE* were higher than the other three exporters (*thrE*, *ncgl0254* and *ncgl0255*). The relationships between the four l-serine exporters were analysed by comparing changes in transcription levels in SAAAI, SSAAIΔ*thrE*, SSAAIΔ*serE*, SSAAIΔ*ncgl0254* and SSAAIΔ*ncgl0255*. The results showed that when one of the exporters was deleted, the transcription levels of the other exporters were increased to compensate and prevent a dramatic loss in l-serine export capacity. As shown in Fig. 5B, when one of the exporters was deleted, the transcription levels of the other exporters were increased. Specifically, when *serE*, *ncgl0254* or *ncgl0255* were deleted, the transcription levels (Log_2_ Fold Change) of *thrE* in the single deletion strains were consistently highest, and when *serE* was deleted the Log_2_ Fold Change reached 8. By contrast, the change in the transcription level of *serE* when one of the other exporters was deleted was negligible. This may be because the expression level of *thrE* was only 2.3% that of *serE* in strain SSAAI, consistent with ThrE exporting l-serine at a low efflux rate (18). Indeed, ThrE may only be used as a supplementary exporter when there are no other more effective l-serine exporters available. Although the transcriptional levels of exporters responded positively to the high intracellular l-serine concentration caused by deletion of one exporter, comparing the l-serine titers of the deletion strains showed that upregulation of transcription levels could not fully recover l-serine production in single deletion strains. When one of the three exporters (*thrE*, *ncgl0254* and *ncgl0255*) was deleted alongside *serE*, the l-serine titer of double deletion strains was decreased by 60%, 66.7% and 66.1%, respectively, compared with SSAAI, a slight difference from SSAAIΔ*serE* (56.6%). However, when the four exporters were deleted together, the l-serine titer was decreased by 79.7%. The results indicate that, when all four exporters were deleted, upregulation of other exporters could not fully compensate for the absence of the four l-serine exporters. These results support the conclusion that the four l-serine exporters undertake most of the l-serine export. In the process of amino acid production, these exporters cooperate with each other, and they are indispensable for full export capacity. Thus, efficient export of l-serine can be achieved by combinatorial optimisation the expression levels of these exporters (39).

Expression of amino acid exporters in *C. glutamicum* is regulated by transcription factors and their effectors (29, 40). Clarifying the regulatory mechanisms of exporters can lay a solid theoretical foundation for further improving the amino acid titer. However, identifying transcription factors is more difficult than identifying amino acid exporters. To date, only four transcription factors involved in amino acid export have been identified in *C. glutamicum*; the transcription factor LysG regulating LysE, related to export of l-lysine and l-arginine; the transcription factor CgmR regulating CgmA, related to export of l-arginine; and the transcription factor Lrp regulating BrnFE, related to export of branched-chain amino acids (13, 14, 41). Recent studies have focused on overexpressing transcription factors regulating amino acid exporters, or using them as biosensors for screening strains with high amino acid production (13, 35, 42). Herein, we identified SerE as the major l-serine exporter, but when SerE was overexpressed in SSAAI, the l-serine titer was only increased by 10.5%. Subsequent studies found that SerE was regulated by the transcription factor NCgl0581. When *ncgl0581* and *serE* were overexpressed together, the l-serine titer was increased 12%, but the strain displayed a lower growth rate (19). These results suggest that the relationships between SerE, NCgl0581 and l-serine need further investigation. We can conclude that the major l-serine exporter SerE needs to be precisely regulated to balance cell growth and l-serine production. Fine regulation of the major exporter and associated transcription factors may significantly increase the amino acid titer in *C. glutamicum*.

## MATERIALS AND METHODS

### Strains, plasmids and growth conditions

The strains and plasmids used in this study are listed in Table 2. *E. coli* JM109 was used as the cloning host, and was grown in lysogeny broth medium (containing 5.0 g/L yeast, 10.0 g/L tryptone, and 10.0 g/L NaCl) at 37°C with shaking at 220 rpm. Strain SYPS-062, capable of producing l-serine, was isolated from soil, with an l-serine titer of 6.6 g/L. The *C. glutamicum* SYPS-062-33a strain obtained by mutation of SYPS-062 had an l-serine titer of 13.2 g/L. The engineered SSAAI (CGMCC No.15170), selected as the original strain, was constructed in our laboratory by deleting 591 bp of the C-terminal domain of *serA*, deleting *sdaA*, *avtA*, and *alaT*, and attenuating *ilvBN* in the genome of *C. glutamicum* SYPS-062-33a (CGMCC No. 8667), and it had an l-serine titer of 26 g/L. Strain A36, derived from SSAAI by ARTP mutation, had an l-serine titer of 31 g/L, higher than that of SSAAI. The seed and fermentation media for *C. glutamicum* were prepared as described previously (24). The *C. glutamicum* strains were preincubated in seed medium overnight to an optical density (OD_562_) of about 25, then inoculated at OD_562_ = 1 into a 250 mL flask containing 25 mL of fermentation medium and cultured at 30°C with shaking at 120 rpm. The antibiotic kanamycin (50 mg/L) was added when necessary. Samples were withdrawn periodically for measuring residual sugar, amino acids, and OD_562_ as described previously (24).

**TABLE 2.**
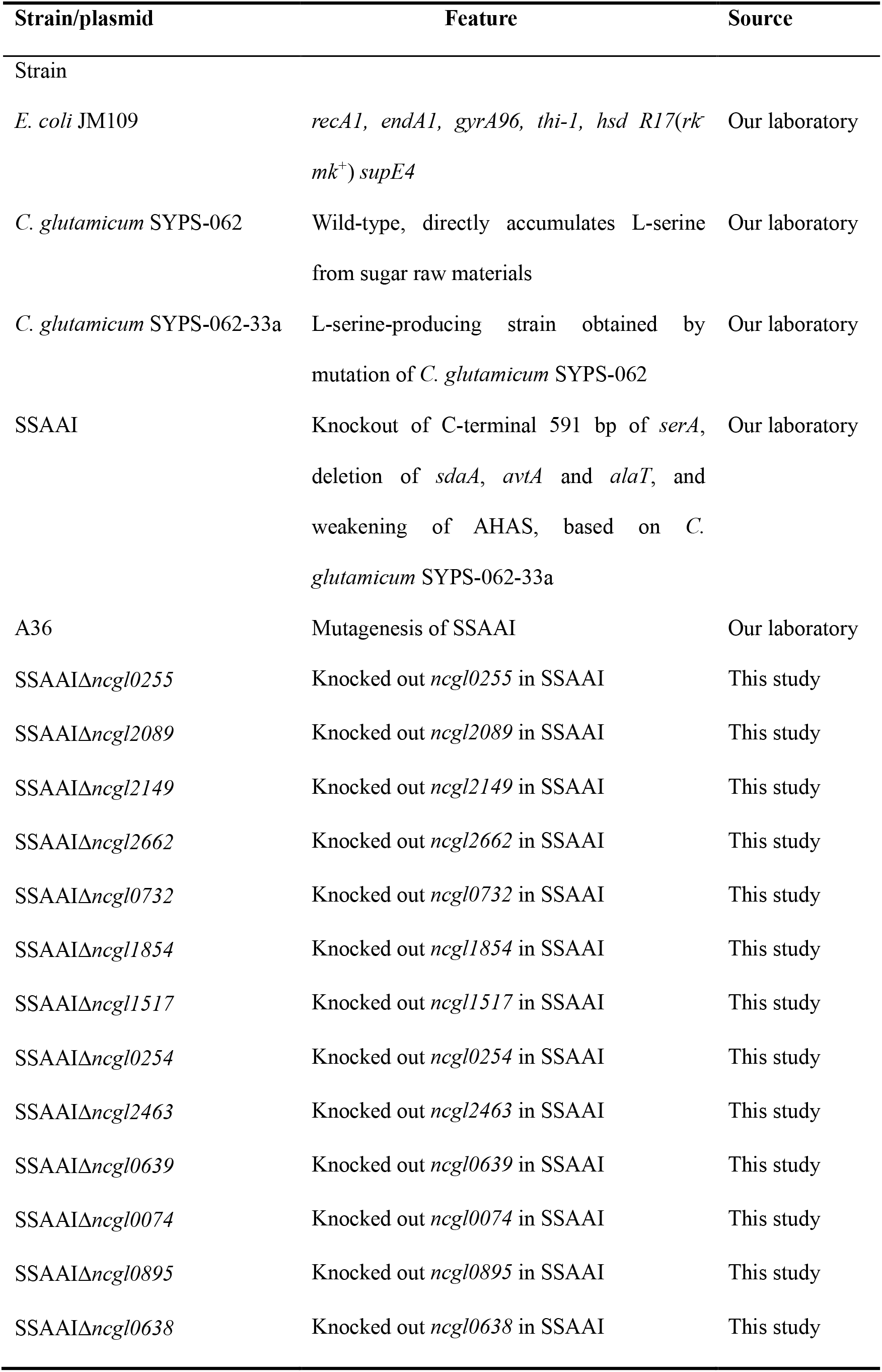
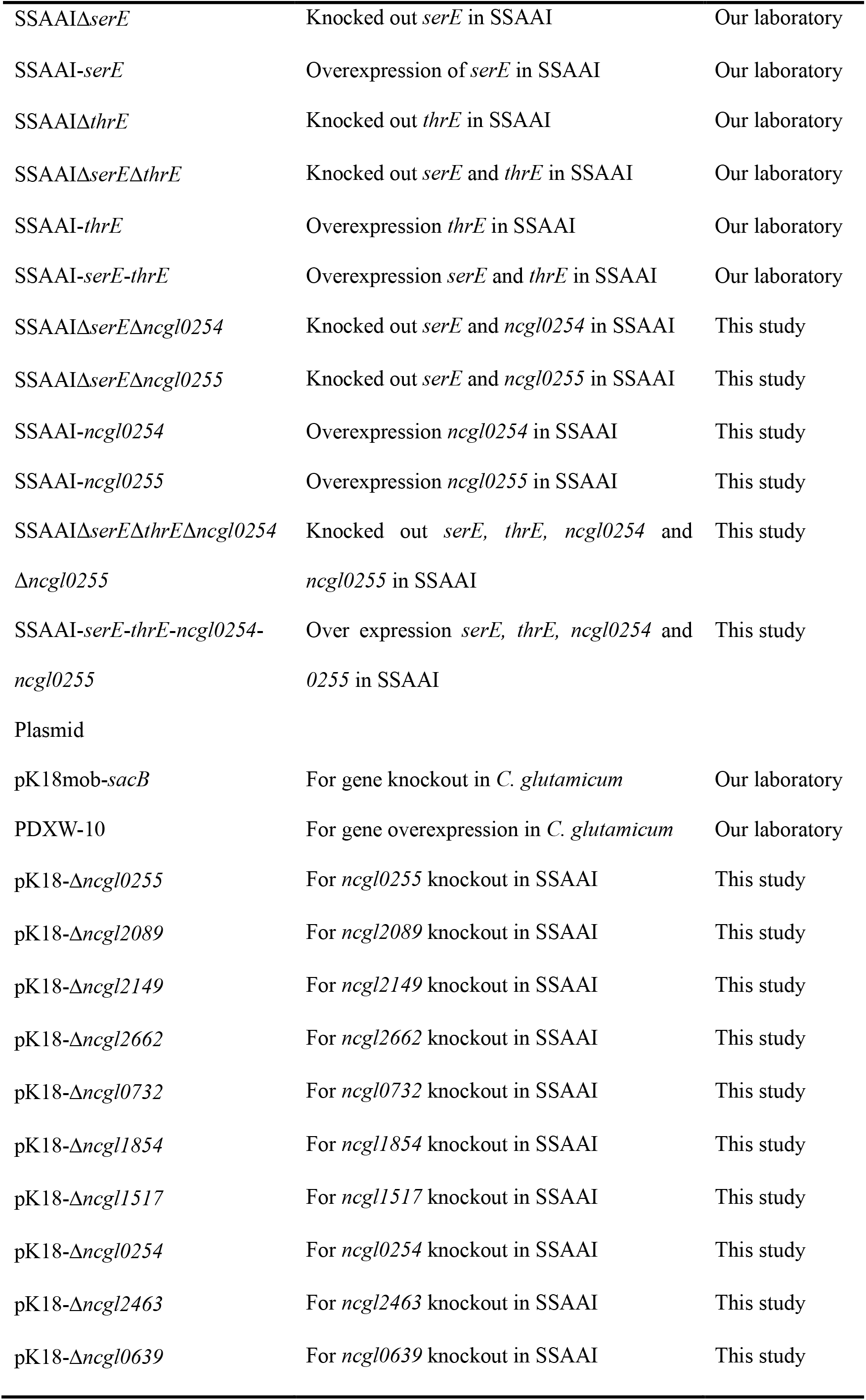
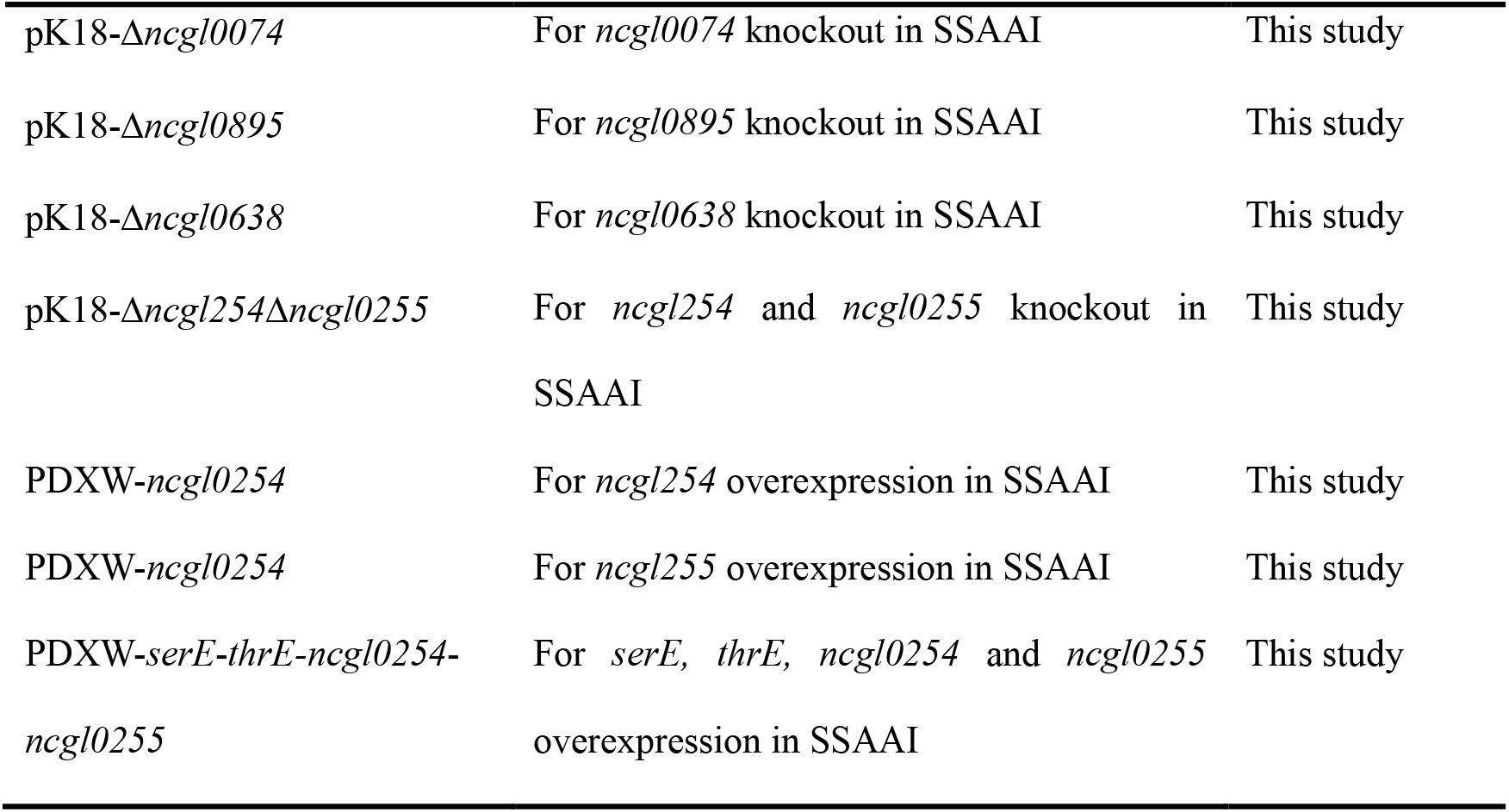
Strains and plasmids used in this study

### Construction of plasmids and strains

The primers used in this study for gene deletion are listed in Table 3. Gene deletion was performed using the nonreplicable deletion vector pK18mob-*sacB* as reported previously (43). For example, to achieve *ncgl0254* deletion, the homologous arm fragments for the *ncgl0254* deletion were amplified from the SSAAI chromosome using primer pairs 0254-1/2 for the upstream fragment and 0254-3/4 for the downstream fragment. Using the two fragments as templates, crossover PCR was performed using primer pair 0254-1/4. The truncated product of *ncgl0254* was digested with *Xba* I and *Hin*d III and ligated to the vector pK18mob-*sacB* treated with the same restriction enzymes. The recombinant plasmid pK-Δ*ncgl0254* was transformed into SSAAI competent cells by electroporation, and chromosomal deletion was performed by selecting cells that resistant to kanamycin but not sucrose, and verified by PCR. The pDXW-10 plasmids were used to overexpress genes in *C. glutamicum* (44). The recombinant plasmids were constructed as follows: the genes *thrE*, *serE*, *ncgl0254* and *ncgl0255* were amplified, digested, and ligated to the pDXW-10 plasmid previously digested with *Hin*d III and *Bgl* II. Strains were generated by electroporation with the corresponding plasmids.

**TABLE 3.**
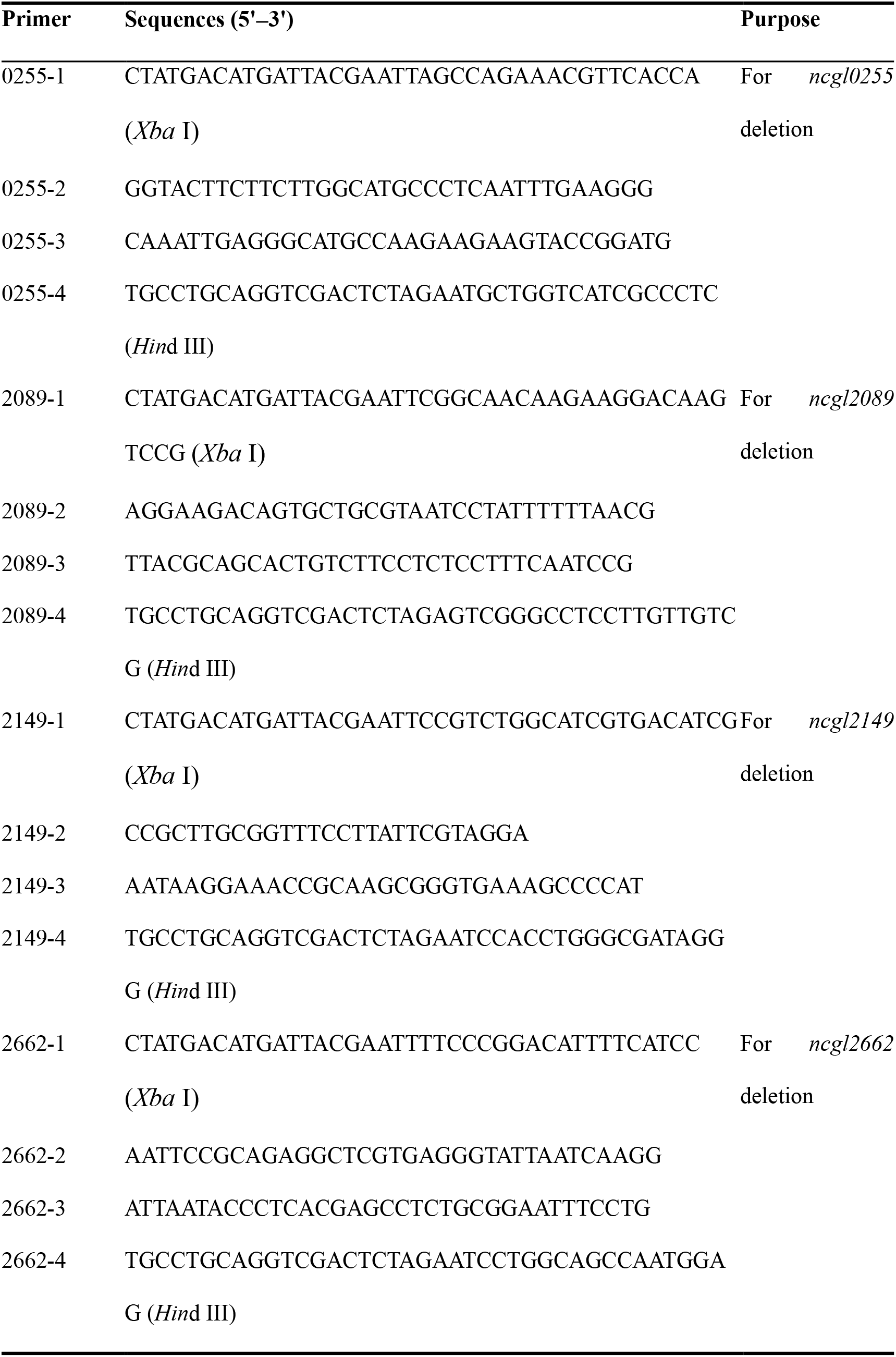
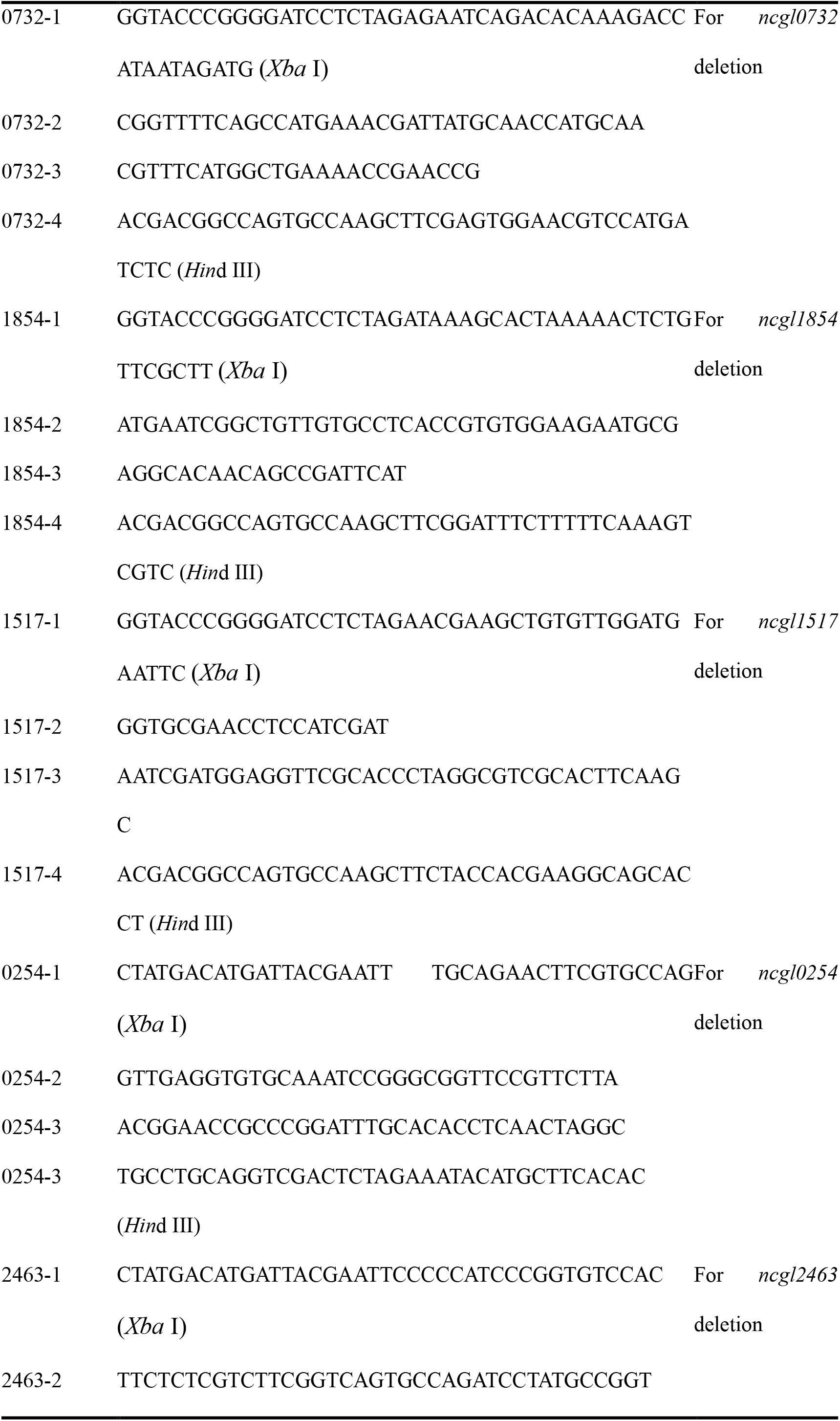
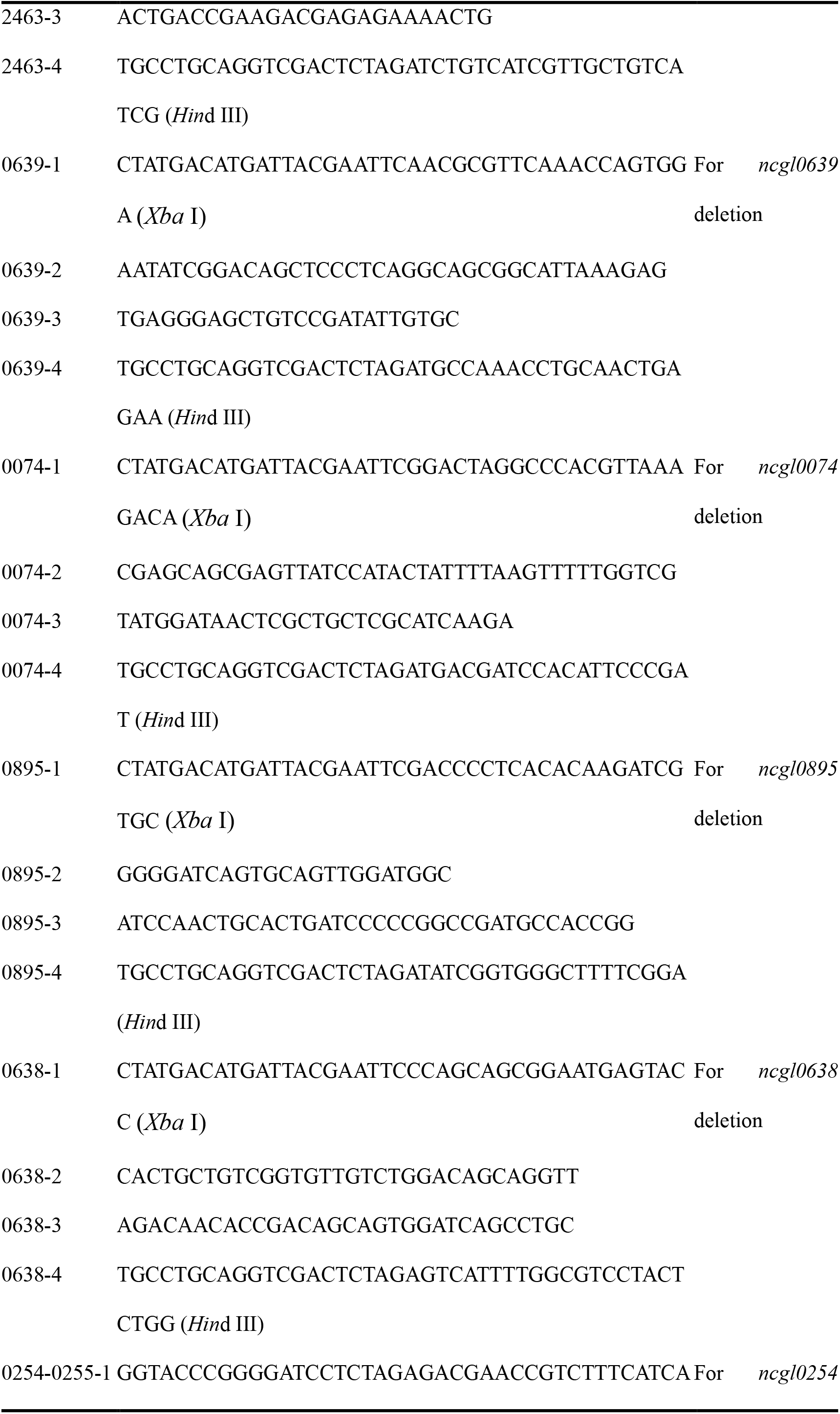
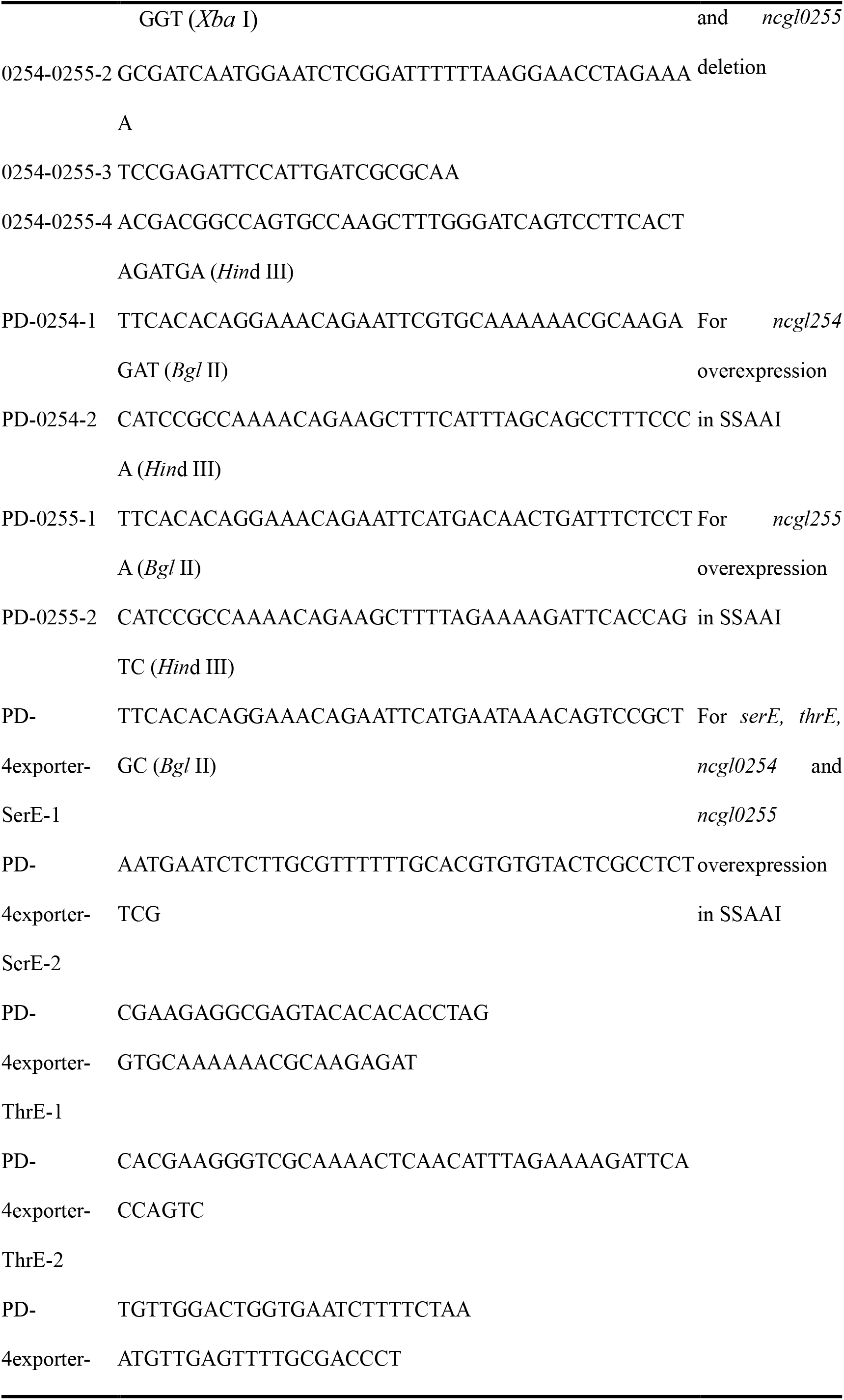
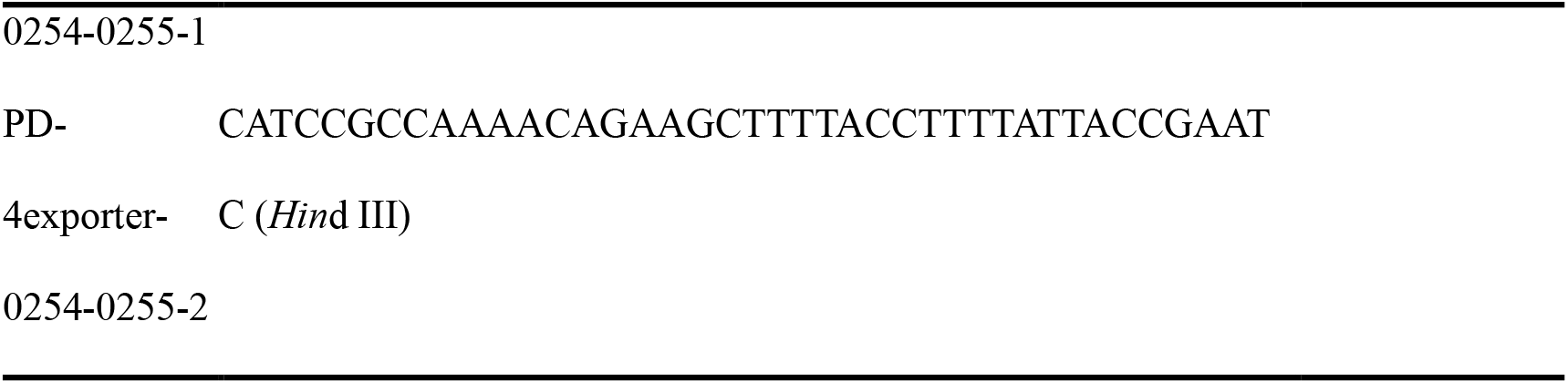
Primers used in this study

### Transcriptome sequencing

The *C. glutamicum* SYPS-062, SYPS-062-33a, SSAAI, and A36 strains were cultured for 60 h then frozen in liquid nitrogen. Transcriptome sequencing and raw data analysis for all four strains were performed by Sangon Biotech (Shanghai, China).

### Amino acid export assay

To verify the functions of NCgl0254 and NCgl0255, a dipeptide Ser-Ser addition assay was performed (18). In brief, cells preincubated in seed medium were washed once with CGXII minimal medium, inoculated into CGXII minimal medium with 2 mM Ser-Ser (dipeptide), and incubated for 2 h at 30°C. Cells were harvested, washed once with cold CGXII minimal medium, and resuspended in CGXII minimal medium (45). Amino acid excretion was initiated by adding 2 mM Ser-Ser. High-performance liquid chromatography (HPLC) was used to determine the titers of amino acids (46).

### Analytical procedures

Cell density (OD_562_) was measured using an AOE UV-1200S UV/vis spectrophotometer (AOE Instruments Co. Inc., Shanghai, China). For measurement of extracellular l-serine titer in shake-flask fermentation, 500 μL of culture was centrifuged at 700 *g* for 5 min, and the supernatant was used for detection after appropriate dilution. The titer of intracellular l-serine was analysed by HPLC using phenyl isothiocyanate as a precolumn derivatisation agent, according to our previous study (24).

### Real-time fluorescence quantitative PCR

*C. glutamicum* bacteria were sampled at different culture time points and total RNA was extracted using a UNlQ-10 Column Trizol Total RNA Isolation Kit (Sangon Biotech). The titer and purity of RNA were determined, and RNA was stored at −80°C until use. Reverse transcription to obtain cDNA was performed according to the instructions of the PrimeScript 1st Strand cDNA Synthesis Kit (Takara Biomedical Technology (Beijing) Co., Ltd.). A ChamQ Universal SYBR qPCR Master Mix Kit (Nanjing Vazyme Biotech Co. Ltd) was used to accomplish quantitative real*-*time PCR with primers listed in Table 4. Experimental data were analysed by GraphPad 8.0.

**TABLE 4.**
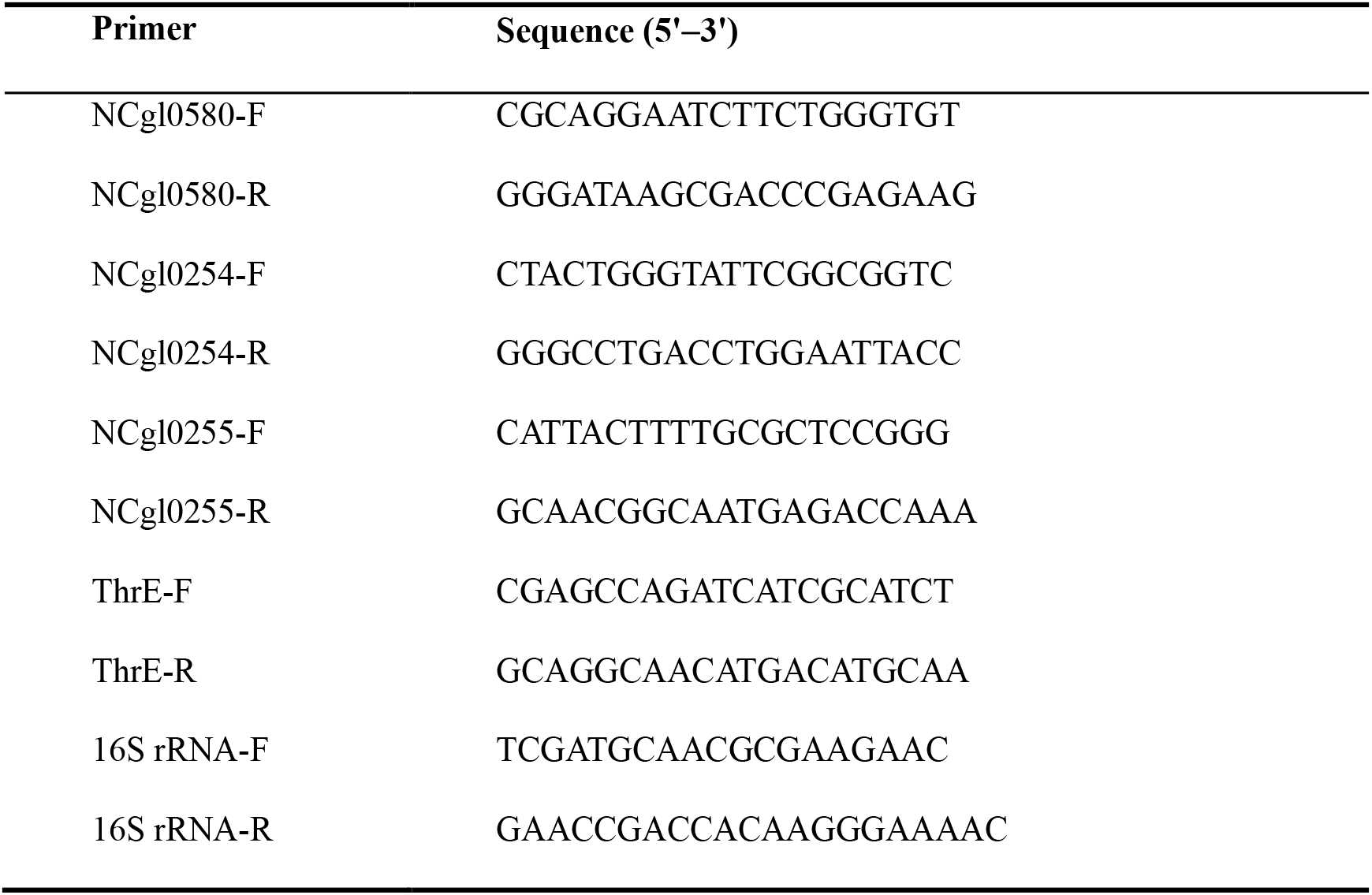
Primers used for quantitative real-time PCR

## ACKNOWLEDGEMENTS

This work was financially supported by the National Key Research and Development Program of China (2018YFA0901400), and the National Natural Science Foundation of China (32171470).

Y.G. and X.Z. designed the research; Y.G. performed experiments; Y.G. and X.Z. analysed the results; Y.G., X.Z. and G.X. wrote the manuscript; Y.G., X.Z., G.X., X.Z., H.L., J.S. and Z.X. edited the manuscript.

We declare no competing financial interests.

